# Redundancy masking and the compression of information in the brain

**DOI:** 10.1101/2025.05.30.657088

**Authors:** Dogukan Nami Oztas, Li L-Miao, Bilge Sayim, Nihan Alp

**Author notes:** Co-first authors (Dogukan Nami Oztas, Li L-Miao). Correspondence concerning this article can be addressed to Nihan Alp. Postal address: Sabanci University, Department of Psychology, Istanbul, Turkey.

## Abstract

The visual world is inherently complex, presenting far more information than a human visual system can process in full. To manage this overload, the visual brain employs several mechanisms. One mechanism that possibly contributes to the reduction of information is redundancy masking (RM): the reduction of the number of perceived items in repeating patterns. For example, when three identical lines are presented in the periphery, observers often perceive only two. The underlying neural mechanisms of RM remain unclear. Here, we use steady-state visual evoked potential (SSVEP) to examine whether redundancy-masked items are neurally suppressed or integrated with neighboring items. Three identical arcs (quarter-circles; 0.44° line width) were presented in the periphery (eccentricities: 17.3°, 19.5°, and 21.7°), each tagged with a unique frequency. Participants maintained central fixation, monitored via a gaze-contingent control, and reported the number of arcs they predominantly perceived after each 10s trial. We analyzed baseline-corrected amplitudes at each tagged frequency and calculated signal-to-noise ratios (SNRs) for fundamental and intermodulation (IM) components, separating trials by behavioral responses (RM: 2 items perceived, non-RM: 3 items perceived). Fundamental frequency comparisons revealed that the outer arc elicited higher SSVEP responses than the inner and middle one under RM, with no significant differences between arcs under non-RM. However, fundamental frequency SNRs did not differ between RM and non-RM perceptions. When we compared IM SNRs, the middle and outer arc’s combination was significantly higher during RM compared to non-RM, suggesting increased neural integration between them. These results indicate that RM involves a loss of conscious access to visual information, yet corresponding neural signals are not entirely suppressed. Instead, the neural signatures we found suggest the integration with neighboring elements across space and time. We suggest that redundancy-masked items – although unavailable for conscious report– are still observed in the neural signatures of RM.

## Introduction

The visual world is inherently complex, with visual objects rarely encountered in isolation. Instead, visual elements are typically embedded within richly cluttered environments, leading to pervasive competition among stimuli for limited perceptual resources. Consequently, the human visual system operates under certain constraints to efficiently prioritize, compress, or omit incoming visual information to manage perceptual overload. One strategy for managing such a complex visual environment is redundancy masking (RM), the phenomenon where observers systematically underreport the number of items in repeating patterns (Sayim & Taylor, 2019; Yildirim et al., 2020). For example, when three aligned identical letter Ts are presented in the visual peripheral, observers often report perceiving only two (Sayim & Taylor, 2019). This highlights a significant omission error, with up to 33% of the visual information (one out of three presented items) being lost, and failing to reach conscious awareness. Investigating RM thus provides crucial insights into how peripheral vision selectively compresses redundant visual information, shedding light on fundamental constraints in visual processing.

Recent studies have revealed several key characteristics of RM. For example, RM strongly depends on item spacing, diminishing as the spacing between repeated elements increases, and completely ceasing at larger spacings (Yildirim et al., 2020). Additionally, spatial arrangement plays a crucial role, with RM being predominantly strong in radially arranged elements but largely absent in tangential configurations (Yildirim et al., 2020). The regularity of the stimulus array modulates RM, as regularly spaced items produce stronger RM effects compared to irregularly arranged patterns (Yildirim et al., 2020). Moreover, item size affects RM, with thinner lines yielding stronger RM than thicker ones (Yildirim et al., 2020). RM also shows atypical visual asymmetries, being stronger on the horizontal than the vertical meridian (Yildirim et al., 2022). However, the neural mechanisms underlying RM remain unclear.

One of the fundamental questions regarding RM is which element within a repeating pattern is predominantly affected and consequently fails to reach conscious perception. Yildirim et al.(2019) addressed this question by presenting arrays of 3-7 identical, equally spaced lines in the visual periphery. Participants first reported how many lines they perceived. Following this task, participants completed a spacing judgment, which differed across two experiments. In Experiment 1, they judged the perceived spacing between adjacent lines; in Experiment 2, they estimated the overall horizontal extent of the array. There were two key findings: First, in RM trials with three presented and two reported lines, the perceived spacing between the two lines was significantly larger than in the presented array. Second, the perceived extent of the array was significantly smaller than in the presented array. This and other studies provide strong evidence that the centrally located line is redundancy-masked. Moreover, they showed that RM goes hand in hand with a compression of visual space (Yildirim et al., 2019; Sayim et al., 2024).

A possible underlying neural mechanism of RM is suppression, which modulates the responses of neurons to competing stimuli, effectively filtering and prioritizing visual information (Baker et al., 2021). In this context, it would mean that the redundancy-masked item(s) are suppressed by neighboring stimuli. Another possible mechanism is integration: following the initial processing of individual elements within the array, redundancy-masked item(s) are (perceptually) integrated with their surrounding components at a later stage of visual processing. Both of these mechanisms are proposed to lead to visual masking—a perceptual phenomenon where the visibility or detectability of a target stimulus is reduced or eliminated by the presentation of another stimulus (Breitmeyer, 2007). However, what mechanism underlies RM remains unclear.

To investigate whether RM involves the suppression of masked (features of) items or their pooling (integration) with neighboring elements, Sayim and colleagues (2024) investigated whether typical masking or pooling effects occurred in RM. Observers viewed arrays of 3-5 radially arranged bars of varying widths (0.1°, 0.25°, 0.4°, 0.55°) presented briefly in the periphery, and participants reported the number of bars they perceived. Following this, a probe with the reported number of bars was displayed. Participants were asked to adjust the widths and spacings of the probe to match the perceived stimulus. The findings revealed that, except for the thinnest bars (0.1°), the perceived thickness of the reported bars did not demonstrate the increase expected if the redundancy-masked bars had been spatially pooled with the perceived bars. If pooling had occurred, the masked item should not have been lost entirely but instead combined with its neighbors, thereby influencing the perceived thickness of the reported bars. This absence of increased widths of the perceived bars indicated that spatial pooling is unlikely to underlie RM, and suggests that redundancy-masked items may be lost, possibly by suppression.

A recent study investigated whether features of redundancy-masked items are integrated with surrounding elements. In particular, Li et al. (2021) presented 3 to 5 concentric circles, and asked participants to report the number of perceived circles. One of the circles contained a unique feature: a small gap located on either the horizontal meridian (in the left or right visual field), or on the vertical meridian (in the upper or lower visual field). Participants reported the location of the gap (up, down, left, or right) and on which ring they perceived the gap. Observers accurately indicated the location of the gap. However, there was a strong trend to report the gap on a more eccentric circle (outward feature migration). Interestingly, when the gap was presented on the central of three circles, it was frequently reported to be on the outer item. This was true for both when participants reported the correct number of rings and when they reported two rings (RM occurred). This result suggests that features of redundancy-masked items may survive RM and be perceived as features of other items, possibly due to integration with more eccentric items.

In a recent work by Hansmann-Roth and colleagues (2025), an appearance-based approach was employed, presenting participants with arrays of lines with varied contrast polarities. The task was to report the perceived number of lines. Subsequently, a probe with the number of lines reported was shown on the screen. Participants were asked to reproduce the perceived contrast polarity pattern by indicating whether each line was black or white. They found that participants largely preserved global features like the stimulus edges and overall contrast polarity in their responses, even when the number of perceived lines was reduced due to RM. These findings indicate that RM occurs after the visual system has grouped and segmented individual items, suggesting that the brain compresses redundant information primarily within these already formed perceptual units. While these studies have advanced our understanding of RM, its neural correlates remain largely unknown. Here, we used EEG and steady-state visually evoked potentials (SSVEPs; Regan, 1966) to investigate RM, as this method allows for the neural tracking of simultaneously presented individual items and their neural interactions.

SSVEPs are continuous oscillatory brain responses elicited by visual stimuli modulated at specific frequencies. We can utilize these SSVEP responses through frequency tagging, a technique where distinct visual stimuli are assigned a flicker frequency. This approach, when applied in EEG, allows us to “tag” and isolate neural signals associated with each individual stimulus. These SSVEP responses are typically observed at two key levels: fundamental frequencies (e.g., f1,f2), which reflect direct neural activity elicited by each individual stimulus flickering at its unique rate, and intermodulation (IM; e.g., f1+f2, f2-f1) frequencies, which arise from non-linear interactions between these stimuli (Gordon et al., 2019). While fundamental frequencies are thought to reflect basic neural responses to visual inputs, IM frequencies often signify integrated processing or complex interactions among those inputs (Norcia et al., 2015).

The ability of SSVEPs to independently track neural signals of multiple stimuli makes them ideal for investigating the underlying neural mechanisms of RM. Specifically, we examined whether the redundancy-masked items were suppressed, which would be reflected as a reduction in fundamental frequency power at the corresponding tag, or whether their neural signals are instead integrated with those of neighboring elements. This integration, indicative of more complex processing, would be evident through the emergence of IM components. For instance, IMs has been shown to reflect feature binding (Aissani et al., 2011), holistic processing (Alp et al., 2016, 2017, 2018), spatial integration in illusory contours (Gundlach & Müller, 2013), and their strength can increase during perceptual competition or when stimuli become perceptually integrated or become invisible, as observed in studies on binocular rivalry or perceptual filling-in (Katyal et al., 2016; Sutoyo & Srinivasan, 2009; Davidson et al., 2020a; 2020b).

Here, we presented 3 identical arcs in the periphery, each flickering at a different frequency (f1 = 4.8 Hz, f2 = 6 Hz, and f3 = 7.5 Hz, allowing us to independently track the neural response to each arc. Participants frequently reported perceiving only two of the three arcs, replicating the classic RM effect (Sayim & Taylor., 2019). We found that all arcs evoked neural responses above noise level, indicated by peaks at their tagged frequencies.This demonstrates that the physical presence of the items still elicited neural activity, even when not consciously reported individually. Interestingly, there were higher fundamental frequency signal-to-noise ratio values for the outer arc compared to the other two arcs when RM occurred. Importantly, among the various IM components observed to be significant above noise levels, specifically f2 + f3 was enhanced during RM, indicating increased neural interaction between the inner and outer items under RM. Our results indicate that RM reflects a loss of conscious access to visual information, but not a complete loss of the neural representation of the masked item(s). Instead, the neural signals associated with redundancy-masked items appear to be integrated with those of neighboring elements across space and time.

## Method

### Participants

A total of 21 students (7 males and 14 females; *M* = 21.19 years, age range: 19–24) from Sabancı University participated in the experiment. Participants were recruited in exchange for course credit. All participants reported normal or corrected-to-normal vision and were naïve to the purpose of the study. Informed consent was obtained from all participants prior to participation. The study was conducted in accordance with the ethical standards of the Declaration of Helsinki and was approved by the Sabancı University Ethics Committee.

### Apparatus and Stimuli

The experiment was conducted on a desktop computer running PsychoPy (v2023.1.1; Pierce et al., 2022), with a screen refresh rate of 120 Hz. The testing room was dimly lit and equipped with a chin rest, a keyboard, and a portable eye tracker (Eyelink 1000 Portable Duo). Participants were seated at a distance of 57 cm from the screen. The stimuli consisted of three grey arcs simultaneously presented on a black background for 10 seconds. The arcs were displayed at varying eccentricities of 17.3°, 19.5°, and 21.7°, either to the right or left of a central fixation point (a black disc, 0.11° in diameter). Each arc’s contrast was modulated by a sine wave at distinct frequencies throughout the 10-second stimulus presentation: 4.8 Hz (f1) for the inner arc, 6 Hz (f2) for the middle arc, and 7.5 Hz (f3) for the outer arc. The contrast of the arcs ranged from 0.5 to 1 (mid-grey to white). The lower bound of 0.5 was chosen to ensure continuous visibility, and to prevent transient invisibility during stimulus presentation.

### Procedure

Before the experiment, participants received instructions outlining the task and the response procedure. To ensure unbiased responses, participants were not informed about the actual number of arcs presented in each trial. Each trial started with a 1-second blank screen displaying only the central fixation point, followed by the three arcs being presented either on the left or right side of the fixation. During this stimulus presentation, participants were instructed to maintain their gaze steadily on the central fixation point and to avoid blinking. A gaze-contingent paradigm was used to track both eyes. Any gaze deviation beyond a 2-degree visual angle from the center, or a blink, triggered an immediate auditory warning, and caused the arcs to disappear until fixation was reestablished.

After each trial, participants reported the number of arcs they predominantly perceived by pressing the corresponding number key on the keyboard. To reduce the physical strain of maintaining fixation without blinking, each trial was followed by a self-paced interval. Participants were allowed to take short breaks as needed and pressed the spacebar to initiate the next trial when ready. The experiment was divided into four blocks, each containing an equal number of left and right visual field presentations. Prior to each block, the eye-tracker was calibrated. A new block was only initiated upon successfully calibrating the eye-tracker, ensuring accurate gaze monitoring throughout the subsequent trials. Breaks between blocks were self-paced. Continuous EEG data were recorded from 64 scalp electrodes positioned according to the International 10/20 System, using an actiCAP (Brain Products) elastic cap. Signals were acquired with BrainVision Recorder (Version 1.24.0001, Brain Products GmbH), and sampled at 1000 Hz.

### Data analysis

#### Behavioral Data

In the behavioral analysis, we examined the number of arcs participants reported. For each trial, the deviation score (DS) was calculated by subtracting the presented number of arcs (i.e., 3) from the reported number of arcs. A negative DS indicates that fewer arcs than presented were reported, a DS of zero represents a correct report of the number of arcs, and a positive DS indicates over-reporting. To assess the occurrence of RM, the individual DS for each trial was averaged, and a one-sample t-test was employed to determine whether the mean DS significantly differed from zero.

#### EEG Data Preprocessing

EEG data were preprocessed using custom MATLAB scripts in conjunction with the EEGLAB toolbox (v2022.0; Delorme & Makeig, 2004). First, raw EEG signals sampled at 1000 Hz were downsampled to 250 Hz to reduce data size and computational load. A bandpass filter (0.1 Hz -50 Hz) was applied to remove low-frequency drifts and high-frequency noise. Electrodes with poor signal quality -identified by amplitudes exceeding typical biological ranges ( ±100 µV), suggesting artifact contamination or poor electrode contact- were identified through visual inspection. These channels were subsequently interpolated using data from their three nearest neighboring channels, a procedure applied to 1.79% of the channels across participants. The data were then re-referenced to the average of all channels to minimize spatial noise. Finally, the EEG data were then segmented into epochs that were time-locked to the stimulus onset, with each epoch spanning the full 10-second stimulus presentation.

#### Spectral Analysis

Frequency-domain analysis was conducted using Fast Fourier Transformation (FFT) to extract steady-state visual evoked potentials (SSVEPs) associated with the frequency-modulated stimuli. Given the 10-second epoch length, the frequency resolution was 0.1 Hz. Prior to spectral analysis, trials were categorized based on behavioral responses (i.e., the number of arcs perceived: 2 or 3 –see Results) and the lateral presentation of stimuli (left or right visual field). This yielded four result conditions. For each participant and channel, trials within each of the result conditions were averaged in the time domain. FFT was then applied to the averaged signal across all EEG channels. Spectral amplitudes were normalized by dividing by N/2 (where N is the number of data points in the epoch) and squared to obtain power values. To calculate the signal-to-noise ratio (SNR) at each frequency, the power at the frequency was divided by the mean power of the second to eighth neighboring frequency bins on both sides of the target frequency, excluding the adjacent bins.

#### Statistical Analysis

Statistical analyses were conducted in R (R Core Team, 2021) using RStudio (RStudio Team, 2015). To determine whether the frequencies of interest were distinguishable from noise in the frequency domain, we first performed time-domain averaging of all trials for each channel and participant. Subsequently, we performed FFT and calculated the SNR, averaging across all channels. The statistical test of whether a frequency is present in the data focused on the fundamental frequencies—f1 = 4.8 Hz, f2 = 6 Hz, and f3 = 7.5 Hz—and their harmonics up to the third order (e.g., for f1; 2×4.8 = 9.6 Hz, 3×4.8 = 14.4 Hz), as well as IM components:f2–f1, f3–f2, f3–f1, f1+f2, f1+f3, f2+f3, and f1+f2+f3 (i.e., 1.2, 1.5, 2.7, 10.8, 12.3, 13.5, and 18.3 Hz respectively). To test whether these frequencies were significantly above the noise level (SNR = 1), we performed one-sample t-tests comparing each target frequency’s SNR against the baseline value of 1. Subsequent analyses were restricted to those fundamental and the IM components that showed SNR values significantly above the noise level.

For statistical comparisons, we used the lme4 package (Bates et al., 2015) to fit linear mixed-effects models (LMMs) and the emmeans package (Lenth, 2025) for post-hoc pairwise comparisons. This analysis exclusively focused on data from the Oz channel, as it represented the only common channel that consistently captured responses across all tagged frequencies. The primary analysis compared SNRs between RM (response 2) trials and correct (non-RM; response 3) trials. Trials in which participants reported 1, 4, and 5 arcs were excluded due to the low number of trials (12.7%, 1.6%, 0.9%, of all trials, respectively). Separate LMMs were constructed for fundamental (4.8, 6.0, 7.5 Hz) and IM frequencies (10.8, 12.3, 13.5, 18.3 Hz) using this single-channel data. In each model, the response number (2 vs. 3) and the frequencies were included as fixed factors. To account for individual differences in SNR, only random intercepts for participants were included. Pairwise comparisons were conducted between response conditions at each specific frequency level, with p-values adjusted using the Tukey method to correct for multiple comparisons.

To analyze differences in frequency-specific SNR among the individual arcs within a response, we ran two additional LMMs on data extracted from predefined regions of interest (ROIs). One model included only the fundamental frequencies, and the other only the IM components. In both models, the number of arcs reported and relevant frequencies were included as fixed factors. Random intercepts were added for both participants and channels nested within participants. Estimated marginal means were used for pairwise comparisons, with Tukey’s method to adjust for multiple comparisons within conditions. Bonferroni corrections were used for cross-condition adjustments.

#### Region of interest

Statistical analyses were conducted using a region-of-interest (ROI) approach, focusing on the channels that exhibited the highest SNR at each set of frequencies of interest. Specifically, we defined two sets of target frequencies: (1) fundamental frequencies and (2) IM frequencies. For each set, we computed the mean SNR for every channel, separately for each hemisphere (left and right presentation). Within each hemisphere, we then selected the top six channels with the highest average SNR to define the ROIs. These selections were made independently for fundamental and IM frequencies (see Results). This approach accounts for the fact that distinct neural processes and generators are often associated with fundamental and IM frequencies. Fundamental and IM frequencies typically show different topographical distributions across the scalp (Boremanse, et al., 2013; Boremanse et al., 2014; Drijvers et al., 2020; Koenig-Robert, et al., 2023). Furthermore, since brain activity related to peripheral visual stimuli is localized in the contralateral hemisphere (e.g., left visual field stimuli activate the right hemisphere), we selected ROI channels separately for each hemisphere to capture possible lateralized processing differences (e.g., Li et al., 2023; Das et al., 2024).

## Results

### Behavioral

#### Deviation Scores (DS)

Participants exhibited a mean deviation score (DS) of -0.76 (SD = 0.41), indicating that, on average, they underreported the number of arcs presented. A one-sample t-test revealed that this deviation was statistically significant, t(20) = -8.37, p < .0001, providing robust evidence for RM that participants perceived fewer items than were displayed.

### EEG

#### Validation of Frequency Components

Prior to the main analysis, we examined which frequency components were distinguishable from noise. The SNR analysis revealed significantly higher values (relative to a baseline of 1) for the fundamental frequencies (4.8, 6.0, 7.5 Hz), specific harmonics (9.6, 12, 15, 18 Hz), and IM frequencies (10.8, 12.3, 13.5, 18.3 Hz). All statistical comparisons reached significance (ps < .05, FDR corrected), indicating the presence of strong frequency-tagged neural responses across participants (Fig. 1).

**Figure 1.**
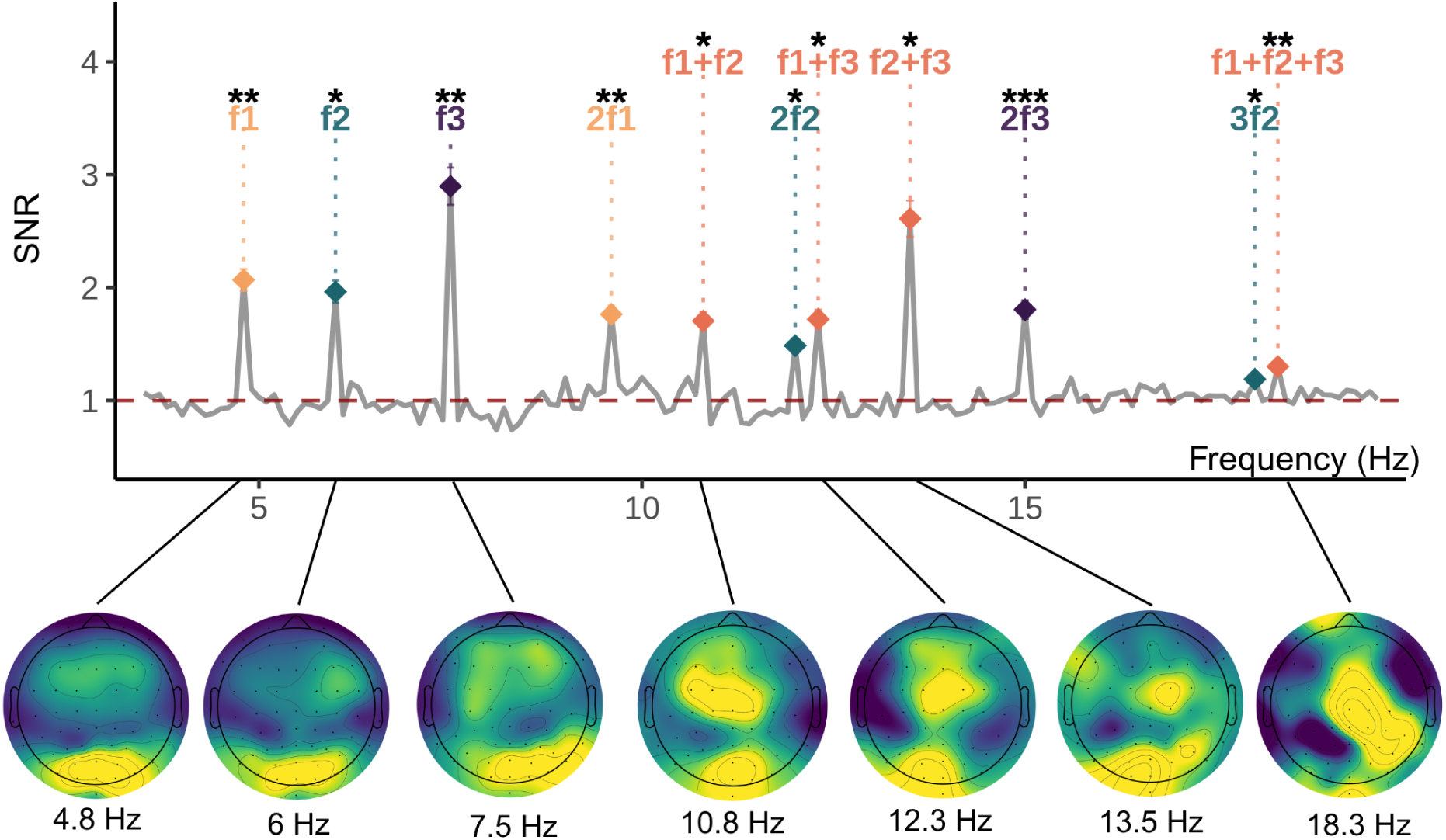
Averaged signal-to-noise ratio (SNR) spectrum from 3 to 20 Hz with a horizontal reference line indicating the noise level ( SNR = 1). Frequencies significantly above the noise level — including the fundamental frequencies, their harmonics, and intermodulation frequencies —are highlighted and labeled. Asterisks indicate that their SNR was significantly greater than 1 (FDR-corrected). The scalp topographies below depict the SNR distributions at each fundamental and IM frequency that are above the noise level.

### Response Comparisons

The analysis of fundamental frequencies revealed no significant differences in SNR between the Response 2 and Response 3 (Fig. 2A) conditions at any of the tested fundamental frequencies (all ps > .05; Fig. 2B). In contrast, the IM frequencies analysis revealed a significant effect at 13.5Hz (f2+f3), where SNR was significantly higher during Response 2 compared to (b = 1.35, t(285.88) = 2.26, p = .025, 95% CI [0.17, 2.53]). No significant differences were observed for the remaining IM frequencies (10.8, 12.3, and 18.3 Hz), all ps > .05 (Fig. 2C).

**Figure 2.**
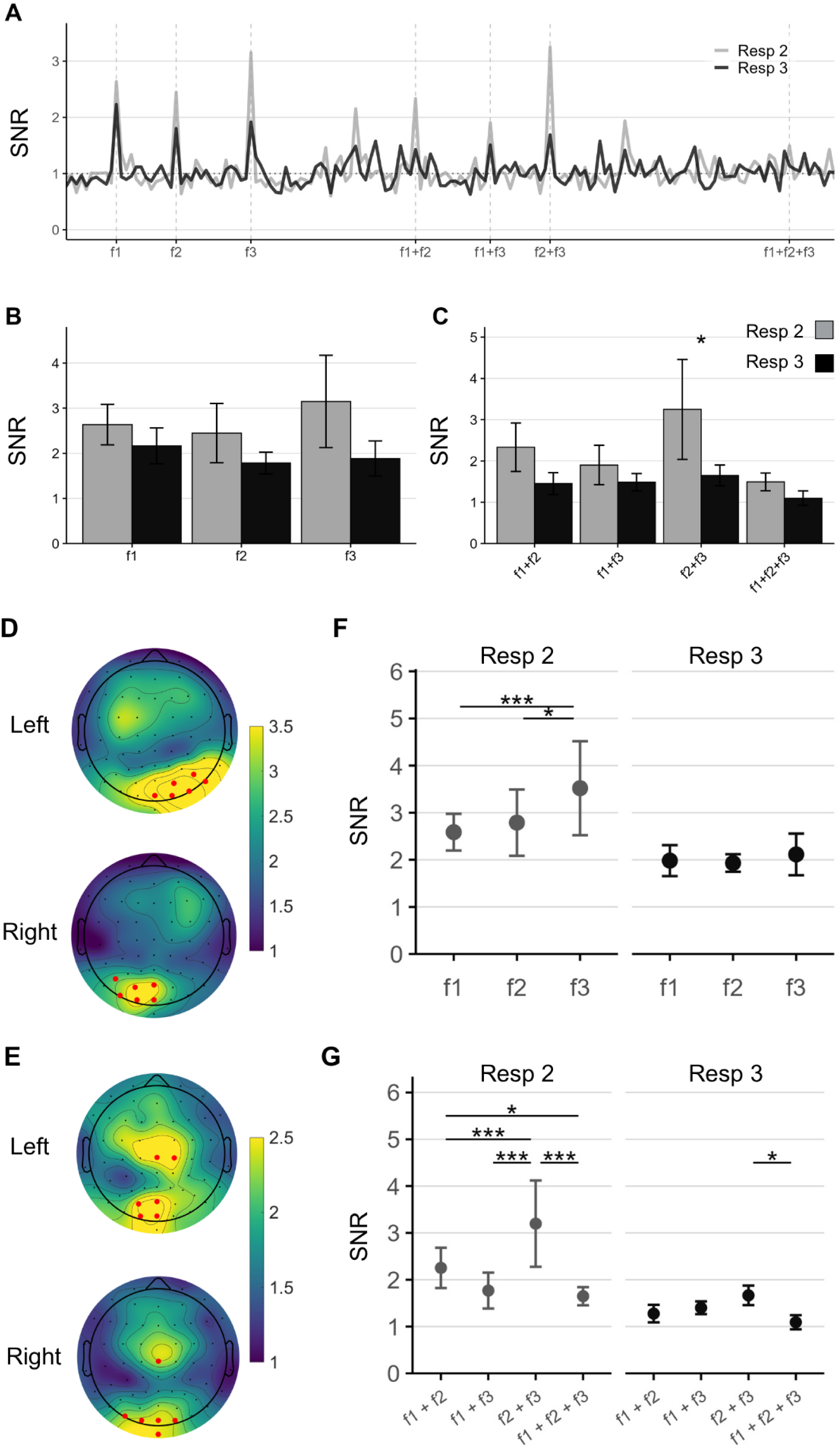
**(A)** SNR spectrum at channel Oz for Response 2 and Response 3 **(B)** Mean SNR (+/- SE) for the fundamental frequencies (f1: 4.8 Hz, f2: 6.0 Hz, f3: 7.5 Hz) plotted separately for the Response 2 and Response 3 conditions at channel Oz. Bars represent the mean SNR across participants; error bars denote standard error of the mean (SE). Asterisks indicate significant differences between response types within a frequency. **(C)** Mean SNR (+/- SE) for intermodulation frequencies (f1+f2: 10.8 Hz, f1+f3: 12.3 Hz, f2+f3: 13.5 Hz, f1+f2+f3: 18.3 Hz) plotted as in panel B **(D)** Scalp topographies illustrating the mean SNR distribution for the fundamental frequencies (4.8, 6, and 7.5 Hz), averaged across participants. Electrodes marked in red represent the ROI channels used for subsequent analyses, shown separately for each hemisphere. **(E)** Mean SNR across all participants at each fundamental frequency, with 95% CI, grouped by the reported number of arcs. Asterisks denote significant pairwise differences. **(F)** Scalp topographies illustrating the mean SNR distribution for the intermodulation frequencies (10.8, 12.3, 13.5, and 18.3 Hz), with selected ROIs highlighted. **(G)** As in panel (E), mean SNR with 95% CI for each intermodulation frequency, separated by the reported number of arcs.

### ROI Selection

All statistical analyses were restricted to an ROI based on channels that showed the highest SNR values at our frequencies of interest. For the fundamental frequencies (4.8, 6, and 7.5 Hz), we calculated the average SNR for each channel and selected the six channels with the highest values in each hemisphere. In the right hemisphere, the selected ROI channels were Oz, O1, POz, PO3, PO7, and P5; in the left hemisphere, they were Oz, O2, PO4, PO8, P6, and P8 (Fig. 2D). The same procedure was used for the IM frequencies (10.8, 12.3, 13.5, and 18.3 Hz). The ROI of the right hemisphere included Oz, POz, O1, PO3, C2, Cz, and Iz, and the left hemisphere ROI comprised Iz, Oz, O1, Cz, O2, and PO7 (Fig. 2E). Only data from these selected channels were included in the subsequent LMMs and post-hoc comparisons.

### Fundamental Frequencies

We fitted an LMM with the reported number of arcs and fundamental frequencies as fixed effects, and random intercepts for participants and channels nested within participants. When participants reported seeing 2 arcs, the outer arc (f3;7.5 Hz) elicited significantly higher SNR compared to the inner arc (f1; 4.8 Hz); b = -0.933, t(1179.17) = -3.688, 95% CI [-1.527, -0.339]) and the middle arc (f2; 6 Hz); b = -0.731, t(1179.17) = -2.888, 95% CI [-1.324, -0.137]. No other significant differences in SNR were observed across frequencies in any condition (*all ps > .05*; Fig. 2F).

### Intermodulation Frequencies

Another model was applied to analyze IM frequencies. When participants reported two arcs, the f2 + f3 component (13.5 Hz) showed significantly higher SNR compared to f1 + f2 (10.8 Hz; b = -0.945, t(1696.92) = -4.704, 95% CI [-1.461, -0.428]), f1 + f3 (12.3 Hz; b = -1.430, t(1696.92) = -7.118, 95% CI [-1.946, -0.913]), and f1 + f2 + f3 (18.3 Hz; b = 1.549, t(1696.92) = 7.713, 95% CI [1.033, 2.066]), indicating a strong non-linear integration between the middle and outer arcs under RM. Additionally, f1+ f3 (12.3 Hz) elicited significantly higher SNR compared to f1 + f2 + f3 (18.3 Hz; b = 0.604, t(1696.92) = 3.009, p = 0.014, 95% CI [0.088, 1.121]). When all three arcs were reported, f2 + f3 (13.5 Hz) elicited higher SNR compared to f1 + f2 + f3(18.3Hz; b = 0.594, t(1696.92) = 2.736, 95% CI [0.036, 1.151]), but no other comparisons among IM components reached significance (all ps > .05; Fig. 2G).

## Discussion

Our study investigated the neural mechanisms underlying redundancy masking (RM) using EEG-based frequency tagging. The behavioral results replicate previous findings (Sayim & Taylor, 2019; Yildirim et al., 2020), showing that participants predominantly perceived fewer items than were presented, most often reporting two instead of three arcs.

We successfully identified detectable neural signals for both fundamental and IM frequencies in our recordings. EEG analyses showed that the neural signals evoked by our stimuli differed significantly depending on whether RM occurred or not. At the fundamental frequencies, SSVEP responses did not differ between the three arcs when all arcs were reported, i.e., when no RM occurred. In contrast, when observers reported two arcs (RM occurred), there were clear SSVP differences between the arcs: the outer arc (f3) elicited significantly stronger responses than the inner and middle arcs. Importantly, fundamental frequency SNRs did not differ between RM and non-RM trials. The analysis of IM components showed significant differences between frequency pairs, particularly when RM occurred. In particular, the f2+f3 component was most prominent, showing significantly higher SNRs during RM compared to no RM, and also being stronger than the other IM components under RM. This result indicates enhanced integration between the middle and outer arcs, suggesting that information from redundancy-masked items is not entirely lost but is likely to be integrated with neighboring items. These findings show that RM is not a result of suppression at early visual stages. Instead, the observed SSVEP response patterns - particularly the enhanced IM responses - show that information from the redundancy-masked items may be integrated with neighboring items during later stages of visual processing.

The pattern of SSVEP responses observed during RM provides important insight into understanding RM. The absence of significant SNR differences between the three arcs during non-RM trials indicates similar neural representations of the stimuli when all items are consciously perceived. However, when RM occurred, the neural signals to the individual arcs exhibited a differential pattern, specifically, the outer arc (f3) showed stronger signals than the other arcs. That is, when RM occurred, the neural responses to the three presented arcs were different, even though the presented stimuli remained the same. Importantly, the persistent SSVEP responses at the fundamental frequencies for all three arcs – even when participants only report two arcs– directly argue against a suppression of the redundancy-masked item at an early stage of visual processing. This implies the physically present but consciously not accessible item still evokes detectable neural signals, demonstrating that visual information is processed but its distinct representations might not reach conscious awareness.

One potential account of the RM is neural suppression, where neural activities and responses to competing stimuli are actively inhibited (Baker et al., 2021). Studies using SSVEPs to investigate various forms of visual masking showed that this kind of neural suppression is known to decrease the fundamental frequency SNRs, filtering out less relevant early visual input (e.g., Vanegas et al., 2025; Tsai et al., 2012; Baker et al., 2021). If RM was due to such neural suppression, we would expect diminished or reduced SNR for the fundamental frequency corresponding to the middle arc. However, our data do not support this idea: there was no significant reduction in fundamental SNR during RM, and all arcs elicited clear SNR responses. Therefore, our results suggest that suppression is unlikely to account for RM.

Instead, our findings support an integration-based account of RM. The key evidence lies in the IM responses, particularly the enhanced f2 + f3 component during RM. IM frequencies, which arise from nonlinear integration between frequency-tagged elements, are widely interpreted as markers of feature integration or binding (e.g., Gordon et al., 2017; Gundlach & Müller, 2013; Alp et al., 2016, 2017, 2018). The selectively increased responses of the f2 +f3 singles - representing the middle and outer arcs - indicate that although the middle arc was not consciously perceived, its neural representation remained, and was likely integrated with the outer arc. This integration was not uniform across all arcs but was specific to the spatial arrangement most affected by RM, i.e., the more eccentric arcs.

This interpretation aligns with previous findings suggesting that features of redundancy-masked items can persist. For example, Li et al. (2021) showed that features of redundancy-masked targets migrated to the outer item, a finding consistent with feature integration. Likewise, Hansmaan-Roth et al.(2025) provide evidence that RM occurs after items have been grouped and segmented, implying that features of the redundancy-masked item persist beyond initial encoding. While their study does not directly demonstrate feature integration, it supports the idea that the redundancy-masked items’ representation is not entirely lost.

Previous research has shown that this feature integration is not based on pooling or low-level averaging. Sayim et al. (2024) directly test this by examining whether the width of identical bars perceived during RM were consistent with predictions from a pooling model. If feature integration was based on such pooling, the perceived items should have spatially integrated the redundancy-masked items, which was not the case. Rather, their results showed that the perceived items were more accurately perceived in RM trials compared to no RM trials. This result demonstrates that RM does not involve simple pooling of features across items. Instead, RM is likely to involve more complex, nonlinear forms of integration - where features from redundancy-masked items persist in the visual system but are not simply averaged with neighboring items. This interpretation is further supported by our IM frequency findings which revealed a selective increased neural signals - particularly in the f2 + f3 responses, indicating integrations between middle and outer items rather than a uniform pooling across all items.

The enhanced IM responses found in our study are also consistent with competition for perceptual dominance. For instance, in perceptual filling-in (PFI), Davidson and colleagues (2020b) observed that IM components exhibited an increase around the time when a target becomes perceptually invisible, which they interpreted as reflecting the competition for perceptual dominance between the target and background elements during the filling-in process. This pattern in IMs is consistent with prior work using binocular rivalry paradigms: Katyal and colleagues (2016) found that IM strength increased during explicit competition for perceptual dominance, peaking just before a switch in awareness of one rivalrous image. Similarly, Sutoyo and Srinivasan (2009) demonstrated that IM responses were sensitive to the perceptual binding of distinct visual inputs (from different eyes) into a coherent percept during binocular rivalry, where these inputs competed for conscious perception. In these studies, IM components reflected dynamic neural interaction between their frequency-tagged target and its context. In our study, the enhanced IM responses between the middle and outer arcs during RM may reflect a similar competition process. That is, the middle arc appears to be the item most vulnerable to RM, possibly due to competitive interactions with its outer neighbor, which is consistent with previous behavioral results that the middle item is the redundancy-masked items (Yildirim et al., 2019; Sayim et al., 2024).

Together, our findings provide neural evidence that RM does not reflect a simple failure of early encoding or a complete suppression of visual information. Instead, RM appears to reflect an integration process where the redundancy-masked item is integrated with outer neighbors during later stages of visual processing. RM leads to a loss of conscious access to the individual identity of the masked item, even though its neural signature persists and selected features may survive masking.

## Acknowledgments

This study was supported by the Scientific and Technological Research Council of Turkey (TUBITAK) under the Grant Number 122N748 and ANR under the Grant Number ANR-19-FRAL-0004. Partial support was also provided by the “PHC Bosphore” programme (project number: 49088TL), funded by the French Ministry for Europe and Foreign Affairs, the French Ministry for Higher Education and Research and the Scientific and Technological Research Council of Turkey.

## CRediT authorship contribution statement

All authors contributed to the study design. Dogukan Nami Oztas collected data. Dogukan Nami Oztas and Li L-Miao analyzed the data. Dogukan Nami Oztas prepared the figures. Dogukan Nami Oztas and Li L-Miao wrote the first draft of the manuscript. All authors interpreted the data and reviewed the manuscript.

## References

Aissani, C., Cottereau, B., Dumas, G., Paradis, A.-L., & Lorenceau, J. (2011). Magnetoencephalographic signatures of visual form and motion binding. Brain Research, 1408, 27–40. 10.1016/j.brainres.2011.05.051.

Alp, N., Kogo, N., Van Belle, G., Wagemans, J., & Rossion, B. (2016). Frequency tagging yields an objective neural signature of Gestalt formation. Brain and Cognition, 104, 15–24. 10.1016/j.bandc.2016.01.008

Alp, N., Kohler, P. J., Kogo, N., Wagemans, J., & Norcia, A. M. (2018). Measuring integration processes in visual symmetry with frequency-tagged EEG. Scientific Reports, 8(1), 6969. 10.1038/s41598-018-24513-w

Alp, N., Nikolaev, A. R., Wagemans, J., & Kogo, N. (2017). EEG frequency tagging dissociates between neural processing of motion synchrony and human quality of multiple point-light dancers. Scientific reports, 7(1), 44012. 10.1038/srep44012

Baker, D. H., Vilidaite, G., & Wade, A. R. (2021). Steady-state measures of visual suppression. PLoS Computational Biology, 17(10), e1009507. 10.1371/journal.pcbi.1009507

Bates D, Mächler M, Bolker B, Walker S (2015). “Fitting Linear Mixed-Effects Models Using lme4.” Journal of Statistical Software, 67(1), 1–48. 10.18637/jss.v067.i01

Boremanse, A., Norcia, A. M., & Rossion, B. (2013). An objective signature for visual binding of face parts in the human brain. Journal of vision, 13(11), 6–6. 10.1167/13.11.6

Boremanse, A., Norcia, A. M., & Rossion, B. (2014). Dissociation of part-based and integrated neural responses to faces by means of electroencephalographic frequency tagging. European Journal of Neuroscience, 40(6), 2987–2997. 10.1111/ejn.12663

Breitmeyer, B. G. (2007). Visual masking: past accomplishments, present status, future developments. Advances in Cognitive Psychology, 3(1-2), 9–20. 10.2478/v10053-008-0010-7

Das, A., Nandi, N., & Ray, S. (2024). Alpha and SSVEP power outperform gamma power in capturing attentional modulation in human EEG. Cerebral Cortex, 34(1), 1–16. 10.1093/cercor/bhad412

Davidson, M. J., Graafsma, I. L., Tsuchiya, N., & Van Boxtel, J. (2020a). A multiple-response frequency-tagging paradigm measures graded changes in consciousness during perceptual filling-in. Neuroscience of consciousness, 2020(1) 10.1093/nc/niaa002

Davidson, M. J., Mithen, W., Hogendoorn, H., Van Boxtel, J. J., & Tsuchiya, N. (2020b). The SSVEP tracks attention, not consciousness, during perceptual filling-in. ELife, 9, e60031. 10.7554/eLife.60031

Delorme, A., & Makeig, S. (2004). EEGLAB: an open source toolbox for analysis of single-trial EEG dynamics including independent component analysis. Journal of neuroscience methods, 134(1), 9–21. 10.1016/j.jneumeth.2003.10.009

Drijvers, L., & Jensen, O. (2020). Rapid invisible frequency tagging reveals nonlinear integration of auditory and visual information. Human Brain Mapping, 42(4), 1138–1152. 10.1002/hbm.25282

Gordon, N., Hohwy, J., Davidson, M. J., van Boxtel, J. J., & Tsuchiya, N. (2019). From intermodulation components to visual perception and cognition-a review. NeuroImage, 199, 480–494. 10.7554/eLife.22749

Gordon, N., Koenig-Robert, R., Tsuchiya, N., Van Boxtel, J. J., & Hohwy, J. (2017). Neural markers of predictive coding under perceptual uncertainty revealed with Hierarchical Frequency Tagging. elife, 6, e22749. 10.7554/eLife.22749

Gundlach, C., & Müller, M. M. (2013). Perception of illusory contours forms intermodulation responses of steady state visual evoked potentials as a neural signature of spatial integration. Biological psychology, 94(1), 55–60. 10.1016/j.biopsycho.2013.04.014

Hansmann-Roth, S., Harmening, W., & Sayim, B. (2025). Compression of visual information in redundancy masking follows grouping and segmentation. Available at 10.2139/ssrn.5254113

Katyal, S., Engel, S. A., He, B., & He, S. (2016). Neurons that detect interocular conflict during binocular rivalry revealed with EEG. Journal of Vision, 16(3), 18–18. 10.1167/16.3.18

Koenig-Robert, R., Pace, T., Pearson, J., & Hohwy, J. (2023, January 10- preprint). Time-resolved hierarchical frequency-tagging reveals markers of predictive processing in the action-perception loop. Authorea Preprint. https://www.authorea.com/doi/full/10.22541/au.167338513.30310833

Lenth R (2025). emmeans: Estimated Marginal Means, aka Least-Squares Means. R package version 1.11.1-00001, https://rvlenth.github.io/emmeans/.

Li, M., Yildirim, F. Z., Alp, N., & Sayim, B. (2021, August). Seeing features of unseen objects: feature migration in redundancy masking. In 43rd European Conference on Visual Perception.

Li, R., Xu, M., You, J., Zhou, X., Meng, J., Xiao, X., Jung, T.-P., & Ming, D. (2023). Modulation of rhythmic visual stimulation on left-right attentional asymmetry. Frontiers in Neuroscience, 17, 1156890. 10.3389/fnins.2023.1156890

Norcia, A. M., Appelbaum, L. G., Ales, J. M., Cottereau, B. R., & Rossion, B. (2015). The steady-state visual evoked potential in vision research: A review. Journal of vision, 15(6), 4–4. 10.1167/15.6.4

Peirce, J. W., Hirst, R. J. & MacAskill, M. R. (2022). Building Experiments in PsychoPy. 2nd Edn London: Sage.

Posit team (2025). RStudio: Integrated Development Environment for R. Posit Software, PBC, Boston, MA. http://www.posit.co/.

R Core Team (2021). R: A language and environment for statistical computing. R Foundation for Statistical Computing, Vienna, Austria. https://www.R-project.org/

Regan, D. (1966). Some characteristics of average steady-state and transient responses evoked by modulated light. Electroencephalography and clinical neurophysiology, 20(3), 238–248. 10.1016/0013-4694(66)90088-5

Sayim, B., & Taylor, H. (2019). Letters lost: Capturing appearance in crowded peripheral vision reveals a new kind of masking. Psychological Science, 30(7), 1082–1086. 10.1177/0956797619847166

Sayim, B., Oztas, D. N., Li, L., & Alp, N. (2024). Seeing less but seeing better: Information loss and accuracy gain in redundancy masking. Journal of Vision, 24(10), 1023–1023. 10.1167/19.10.13c

Sutoyo, D., & Srinivasan, R. (2009). Nonlinear SSVEP responses are sensitive to the perceptual binding of visual hemifields during conventional ‘eye’ rivalry and interocular ‘percept’ rivalry. Brain research, 1251, 245–255. 10.1016/j.brainres.2008.09.086

Tsai, J. J., Wade, A. R., & Norcia, A. M. (2012). Dynamics of normalization underlying masking in human visual cortex. Journal of Neuroscience, 32(8), 2783–2789. 10.1523/JNEUROSCI.4485-11.2012

Vanegas, M. I., Blangero, A., & Kelly, S. P. (2015). Electrophysiological indices of surround suppression in humans. Journal of neurophysiology, 113(4), 1100–1109. 10.1152/jn.00774.2014

Van Overwalle, J., Van der Donck, S., Van de Cruys, S., Boets, B., & Wagemans, J. (2024). Assessing spontaneous categorical processing of visual shapes via frequency-tagging EEG. The Journal of Neuroscience, 44(16), e1346232024. 10.1523/JNEUROSCI.1346-23.2024

Yildirim, F. Z., & Sayim, B. (2022). High confidence and low accuracy in redundancy masking. Consciousness and cognition, 102, 103349. 10.1016/j.concog.2022.103349

Yildirim, F. Z., Coates, D. R., & Sayim, B. (2019). Lost lines in warped space: Evidence for spatial compression in crowded displays. Journal of Vision, 19(10), 13c–13c. 10.1167/19.10.13c

Yildirim, F. Z., Coates, D. R., & Sayim, B. (2020). Redundancy masking: The loss of repeated items in crowded peripheral vision. Journal of Vision, 20(4), 14. 10.1167/jov.20.4.14

Yildirim, F. Z., Coates, D. R., & Sayim, B. (2022). Atypical visual field asymmetries in redundancy masking. Journal of Vision, 22(5), 4. 10.1167/jov.22.5.4

